# Cytoplasmic short linear motifs in ACE2 and integrin β3 link SARS-CoV-2 host cell receptors to endocytosis and autophagy

**DOI:** 10.1101/2020.10.06.327742

**Authors:** Johanna Kliche, Muhammad Ali, Ylva Ivarsson

**Affiliations:** Department of Chemistry, BMC, Uppsala University, Husargatan 3, 751 23 Uppsala, Sweden

**Author notes:** Communicating author.

**Keywords:** SARS-CoV-2 receptors, SLiMs, endocytosis, autophagy, ACE2, integrins, LIR, phospho-regulation, protein-protein interactions

## Abstract

The spike protein of the SARS-CoV-2 interacts with angiotensin converting enzyme 2 (ACE2) and enters the host cell by receptor-mediated endocytosis. Concomitantly, evidence is pointing to the involvement of additional host cell receptors, such as integrins. The cytoplasmic tails of ACE2 and integrin β3 contain a plethora of predicted binding motifs. Here, we confirm the functionality of some of these motifs through affinity measurements. The class I PDZ binding motif in the ACE2 cytoplasmic tail binds the first PDZ domain of the scaffold protein NHERF3. The clathrin-adaptor subunit AP2 μ2 interacts with an endocytic motif in the ACE2 with low affinity and the interaction is abolished by phosphorylation of Tyr781. Furthermore, the C-terminal region of integrin b3 contains a LC3-interacting region, and its interaction with ATG8 domains is enhanced by phosphorylation. Together, our data provides possible molecular links between host cell receptors and endocytosis and autophagy.

**One sentence summary:** Affinity measurements confirmed binding of short linear motifs in the cytoplasmic tails of ACE2 and integrin β3, thereby linking the receptors to endocytosis and autophagy.

## Introduction

A novel coronavirus, severe acute respiratory syndrome coronavirus 2 (SARS-CoV-2), was identified as the pathologic agent for an outbreak of pneumonia in Wuhan, China in December 2019 (1). The virus, which is causing the new lung disease termed coronavirus disease 19 (COVID-19), has infected 30.6 million people and claimed over 945,000 fatalities (09/2020). It is hence posing an immediate health and socioeconomic threat with worldwide impact (2). Understanding how the virus infects and replicates within the host cell is crucial for the development of novel targeted therapies.

SARS-CoV-2 employs the angiotensin converting enzyme 2 (ACE2) as a human host receptor for cell attachment and subsequent receptor-mediated endocytosis (3,4). Structural validation of the interaction has been provided by co-crystallisation of the receptor binding domain of the SARS-CoV-2 spike protein with the ACE2 ectodomain, as well as a cryo-EM structure with full-length ACE2 (5,6). However, analysis of *ACE2* expression across different human tissues revealed surprisingly poor expression in the lungs as compared to, for instance, high expression in the kidney and heart (7,8). The extreme damage that the virus can inflict on infected lungs indicates the requirement to use alternative receptors for host cell attachment and entry. In line, neuropilin-1, abundantly expressed in the olfactory epithelium, was shown to enhance SARS-CoV-2 infection and confirmed as receptor (9). The role of C-type lectins as potential interactors for the SARS-CoV-2 spike protein on innate immune cells has additionally been pointed out (10). Last, integrins have been proposed as candidate receptors due to the presence of a known integrin-binding RGD motif in the spike protein (11). In support, integrin mediated endocytosis is used by a variety of viruses for cell attachment as well as cell entry (12,13). While experimental validation is still awaiting, the structural feasibility of an ACE2-integrin interaction has been analysed (14). In addition, integrin tissue distribution matches SARS-CoV-2 tropism more closely, which includes functional importance of integrin β_1_ expression on type 2 alveolar epithelial cells (15). Furthermore, there are as of now two case reports of COVID-19 patients with concomitant multiple sclerosis background, who were treated with Natalizumab, an antibody effectively blocking integrin α4 binding (16,17). Both reports support a positive correlation between Natalizumab treatment and mild COVID-19 progression.

Considerable research has been focused on the interaction between the SARS-CoV-2 spike protein and cell surface receptor proteins, particularly ACE2 (3,4,9–11). Further insight into the viral entry mechanism may be provided by investigating the downstream effects of the binding event. Taking a bioinformatical approach, Mészàros *et al*. (14) analysed the cytoplasmic tails of ACE2 and integrins and predicted them to contain a number of short linear motifs (SLiMs) that may link host receptor binding to endocytosis and autophagy. SLiMs are typically 3-10 amino acids long stretches found in the intrinsically disordered regions of the proteome. They may serve as binding sites and engage in dynamic proteinprotein interactions (18). Well-known binding motifs include the class I PDZ (PSD-95/Dlg/ZO-1) binding motif (x[ST]xΦ-COO-, where x indicates any amino acid and Φ indicates a hydrophobic amino acid) that directs PDZ domain containing proteins to the C-terminal tails of targets (19), the AP2 μ2 binding endocytic motif (YxxΦ) (20), and the LC3-interacting region (LIR) motif, which mediates the interaction with the ATG8 domain containing proteins MAP1LC3s and GABARAPs in the phagophore membrane (21). Hence, SLiMs on the cytosolic side of the host receptor proteins may allow the recruitment of proteins that enable and mediate receptor and concomitant virus particle internalisation.

Mészàros *et al*. predicted ACE2 to contain binding motifs for AP2 μ2, NCK Src Homology 2 (SH2) domains as well as PDZ and phosphotyrosine binding (PTB) domains (Figure 1). These binding motifs, if functional, would act as molecular platforms linking the receptor to the various biological processes associated with the interacting protein domains. AP2 μ2, for example, is part of the clathrin-adaptor complex formed during clathrin-dependent endocytosis (22). Further, SH2 domains are known phospho-Tyr binders and NCK proteins are specifically involved in the organisation of the actin cytoskeleton (23).

**Figure 1:**
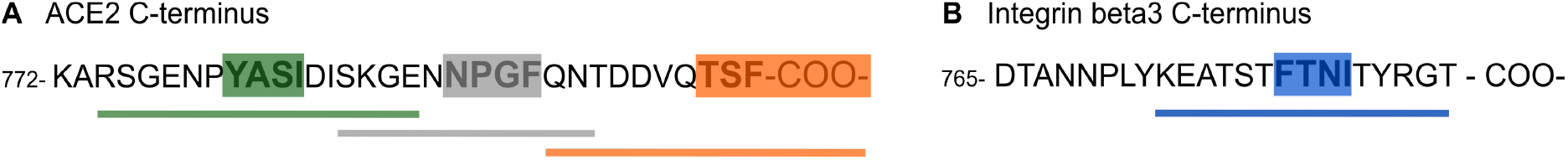
Peptide sequences from the cytoplasmic tails from **A**: ACE2 and **B**: integrin β3. green: region containing predicted overlapping binding sites for NCK SH2 domain, the ATG8 domains of MAP1LC3As and GABARAPs and AP2 μ2; grey: predicted PTB binding site; orange: predicted class I PDZ binding site; blue: predicted ATG8 binding site in integrin β3.

PDZ and PTB domains, on the other hand, are well-established scaffolding domains implicated in the assembly of protein networks (24,25). In addition, Mészàros *et al*. proposed the presence of a phospho-modulated LIR autophagy motif in both ACE2 and the integrin β3 tail (14). If functional, such a motif would provide a link between the receptors, as well as the potential viral entry route to the endocytic-lysosomal pathway.

Phosphorylation of Ser/Thr/Tyr residues may enable, disable or tune a given interaction (26,27). As elaborated by Mészàros *et al*., Tyr781 in the ACE2 tail has emerged as a reproducible phosphosite in high-throughput phosphoproteomics studies (14). In the case of the integrin β tails, it is well-established that phosphorylation regulates their interactions with adaptor and scaffold proteins, for instance allowing competitive binding of PTB domains (28). A bimolecular switch has been suggested in integrin β3, so that phosphorylation of Tyr773 enables binding of DOK1 PTB, whereas phosphorylation of Thr777 creates a 14-3-3 ζ binding site (29). Integrin β tails have moreover a Ser/Thr-rich sequence, that precedes the predicted LIR autophagy motif. The stretch has been proposed to be phosphorylated by Akt and PDK1 (28,30), which would likely modulate the proposed ATG8 domain binding. As rewiring of phospho-signalling has been found as one of the primary host responses to SARS-CoV-2 infection (31), it is important to explore the phospho-regulation of proteinprotein interactions in relation to viral entry and hijacking of the cellular machinery.

In this study we set out to test predicted binding motifs in ACE2 and integrin β_3_ (14) and the potential phospho-regulation of the interactions by affinity measurements.

To this end, we expressed and purified 18 protein domains, and determined affinities with synthetic peptides representing relevant cytoplasmic regions of the receptor tails. We find that the NHERF3 PDZ1 and SHANK1 PDZ domains bind to ACE2 C-terminus. We further confirm that AP2 μ2 binds the cytoplasmic region of ACE2 with low affinity and that the interaction is negatively affected by phosphorylation. Unfortunately, we could not evaluate the functionality of the proposed ACE2 LIR to its full extend due to technical issues. We confirm that the LIR in integrin β3 binds the ATG8 domains of MAP1LC3s and GABARAPs and that the strength of the interaction is increased by ligand phosphorylation.

Taken together, we established the functionality of a subset of the bioinformatically predicted binding motifs, and shed light on the intracellular molecular recognition events that ACE2 and integrin β_3_ engage in, which in a further perspective may contribute to the understanding of how SARS-CoV-2 enters the host cell. On a more general note, identification of a functional phospho-dependent LIR motif in the tail of integrin β_3_ allows to critically link integrin signalling to autophagy.

## Results

We aimed to test the predicted SLiMs in the cytoplasmic tails of ACE2 and integrin β3 for PDZ domains, PTB domains, the μ2 subunit from the clathrin-adaptor AP2 as well as ATG8 domains. We expressed and purified a representative collection of protein domains and used them to explore the affinities for synthetic peptides representing the proposed binding motifs (Figure 1). Affinities were determined by fluorescence polarisation (FP) using fluorescein (FITC)-labelled peptides as direct binders (saturation curves; K_D_ values) or reporters of binding (displacement curves; K_I_ values).

### NHERF3 PDZ1 and SHANK1 PDZ domains bind the ACE2 C-terminus

The presence of a putative classical class I PDZ binding motif in the cytoplasmic tail of ACE2, prompted us to assess direct binding of the C-terminal region to a selection of seven PDZ domains (MAGI PDZ1, NHERF3 PDZ1 and PDZ2, SCRIB PDZ1, SHANK1 PDZ, SNTA1 PDZ and TAX1BP3 PDZ known to interact with class I PDZ binding motif (32), and found in proteins expressed in the lung based on Uniprot annotations.

The labelled C-terminal peptide of ACE2 (FITC-QNTDDVQTSF-COO-) was titrated with increasing concentrations of PDZ domains (Figure 2A). The change in FP signal upon ligand binding was measured and plotted against the protein concentration. Among the seven tested domains, five PDZ domains were classified as non-binders or low-affinity binders of ACE2 as there was no saturation within the tested concentration intervals (0-100 μM or 0-200 μM depending on protein domain). In contrast, the first PDZ domain of NHERF3 and SHANK1 PDZ domain were found to bind with relevant affinities (K_D_ values of 8 and 14 μM respectively; Figure 2A, Table 1).

**Figure 2:**
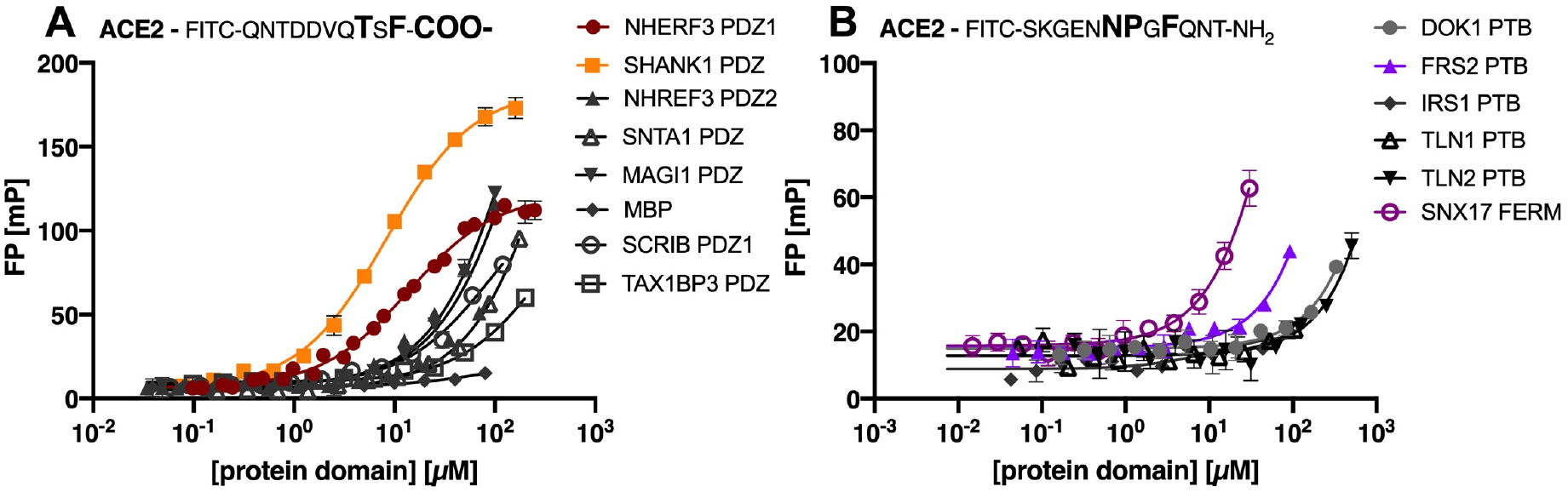
Saturation curves obtained by FP experiments for (**A):** selected class I PDZ domains and (**B)**: a selection of PTB domains and SNX17 FERM. Curves were obtained by titrating the protein domains against the respective FITC-labelled ACE2 peptide, containing either a class I PDZ or a predicted PTB binding motif.

**Table 1:**
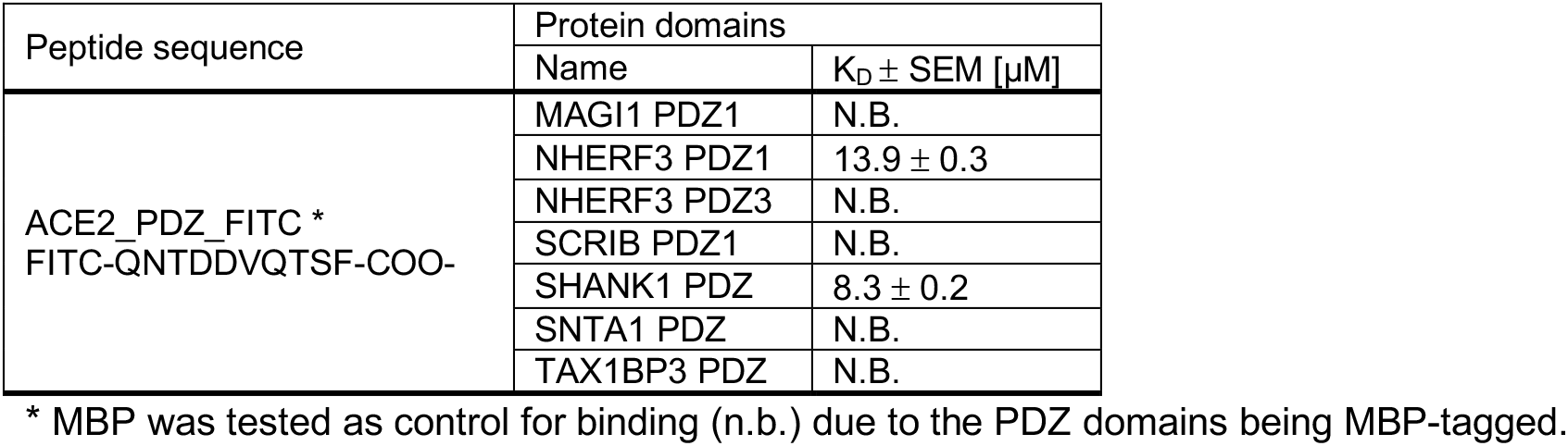
Equilibrium dissociation constant (K_D_) values of the tested PDZ domains for FITC-labelled ACE2 peptides. Indicated errors are the errors of the mean (SEM). N.B. indicates no or low affinity (K_D_ >100 μM).

| Peptide sequence | Protein domains |  |
| --- | --- | --- |
| | Name | $K_D \pm \text{SEM} [\mu\text{M}]$ |
| ACE2_PDZ_FITC *<br>FITC-QNTDDVQTSF-COO- | MAGI1 PDZ1 | N.B. |
| | NHERF3 PDZ1 | $13.9 \pm 0.3$ |
|  | NHERF3 PDZ3 | N.B. |
|  | SCRIB PDZ1 | N.B. |
| | SHANK1 PDZ | $8.3 \pm 0.2$ |
|  | SNTA1 PDZ | N.B. |
|  | TAX1BP3 PDZ | N.B. |

\* MBP was tested as control for binding (n.b.) due to the PDZ domains being MBP-tagged.

### The putative ACE2 PTB binding motif binds to the SNX17 FERM domain with low affinity

We next tested the interactions of the predicted PTB binding motif (NPxF) in the ACE2 C-terminal tail. We attempted to obtain saturation curves for the FITC-labelled ACE2 peptide containing the predicted motif (FITC-SKGENNPGFQNT-NH_2_) and the PTB domains from three proteins (DOK1, IRS1 and FRS2), the PTB-like lobe of the FERM domains of TLN1 and TLN2, and the full FERM domain from SNX17 (Figure 2B). However, none of the tested domains were confirmed as high affinity binders of the ACE2 peptide (Figure 2B). The negative result with the DOK1 PTB domain is in agreement with the bioinformatic analysis, that suggested that the motif should be recognized by a different type of PTB domain (14). Given that there approximately 60 human proteins contain PTB domains (25), a systematic study of PTB domain specificities is needed for a comprehensive picture.

The most promising result was found for the interaction between the SNX17 FERM domain and the ACE2 peptide (Figure 2B), for which we estimate the K_D_ of to be in the range of 200-300 μM. The experiment was however hampered by the low solubility of the protein domain used. In line with this finding, we note that the SNX17 FERM domain bind to NPx(F/Y) containing peptides and has been crystallised in complex with the NPxF containing peptide of KRIT1 (33, 34). The ACE2 peptide thus matches its binding preference. An interaction between the SNX17 FERM domain appears reasonable given that the protein is involved in endosomal recycling of various proteins, including integrins (33,35).

### The ACE2 tail harbours a weak phospho-regulated AP2 μ2 binding motif

The ACE2 tail contains a nine amino acid stretch that is predicted to contain overlapping binding motifs for AP2 μ2, NCK SH2 and the ATG8 proteins (14). In addition, the region contains two phosphosites predicted to serve as bimolecular switches. We aimed at testing the interactions and the predicted phospho-switches in ACE2 binding by measuring the affinities of AP2 μ2, NCK1 SH2 and four ATG8 domains (MAP1LC3A, -B and -C, GABARAPL2) for unphosphorylated and phosphorylated (pTyr781 and pSer783) ACE2 peptides through peptide displacement experiments. Unfortunately, we were not able to determine the affinities for the pSer783 ACE2 peptide due to presence of contaminations that hindered accurate measurement despite ordering the peptide from two different companies.

First, saturation curves for the different protein domains with respective FITC-labelled probe peptides were obtained (Figure 3A, Table 2) to determine appropriate protein domain concentrations for displacement experiments. The labelled peptides for AP2 μ2 (ATG9A) and the ATG8 (SQSTM1) domains were chosen based on our unpublished proteomic peptide phage display data whereas the FITC-labelled peptide (Tir10) for NCK1 SH2 is a reported high-affinity binder (36). Subsequently, ACE2 peptides were titrated against the varying complexes of FITC-labelled peptide and protein domain (Figure 3B), from which IC50 values were derived and K_I_ values could be calculated. However, the ATG8 domains bound neither the unphosphorylated ACE2 nor pTyr781 peptide with relevant affinities (Figure 3B, estimated K_I_ values in mM range). In light of the fact, that the pSer783 peptide binding could not be assessed in the present study, we cannot rule out that the phosphosite interspersed in the predicted LIR motif may enhance affinity, which necessitates an alternative approach (e.g. direct binding with FITC-labelled ACE2 peptide).

**Figure 3:**
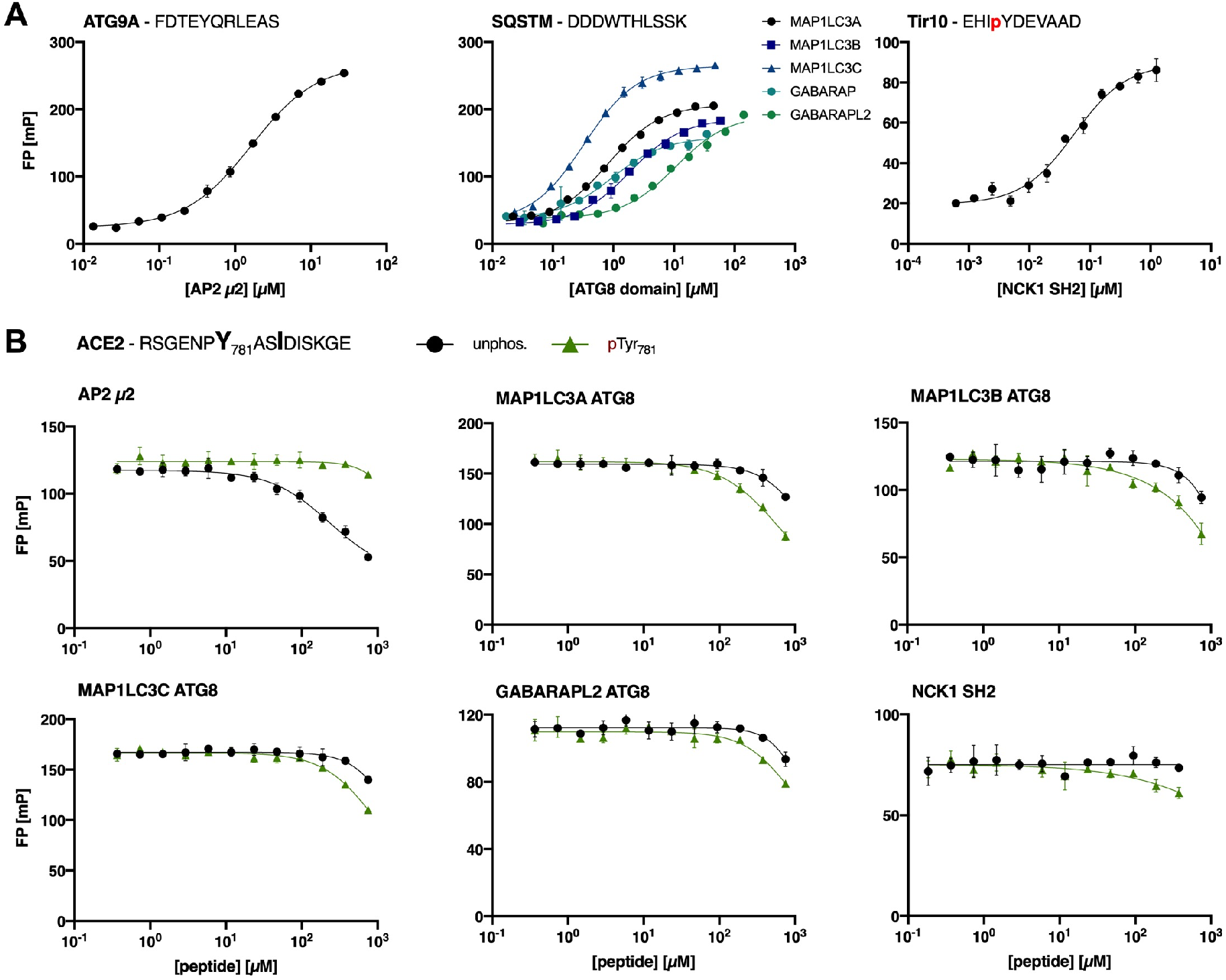
**A**: Saturation curves of AP2 μ2, the ATG8 domains and NCK1 SH2 for the respective FITC-labelled peptides (ATG9A, SQSTM1 and Tir10). **B**: Displacement curves of FP experiments using a peptide from the cytoplasmic ACE2 tail predicted to contain AP2 μ2, ATG8 and NCK SH2 domain binding motifs. Preferential binding of the different domains for unphosphorylated and pTyr781 ACE2 peptide was tested.

**Table 2:**
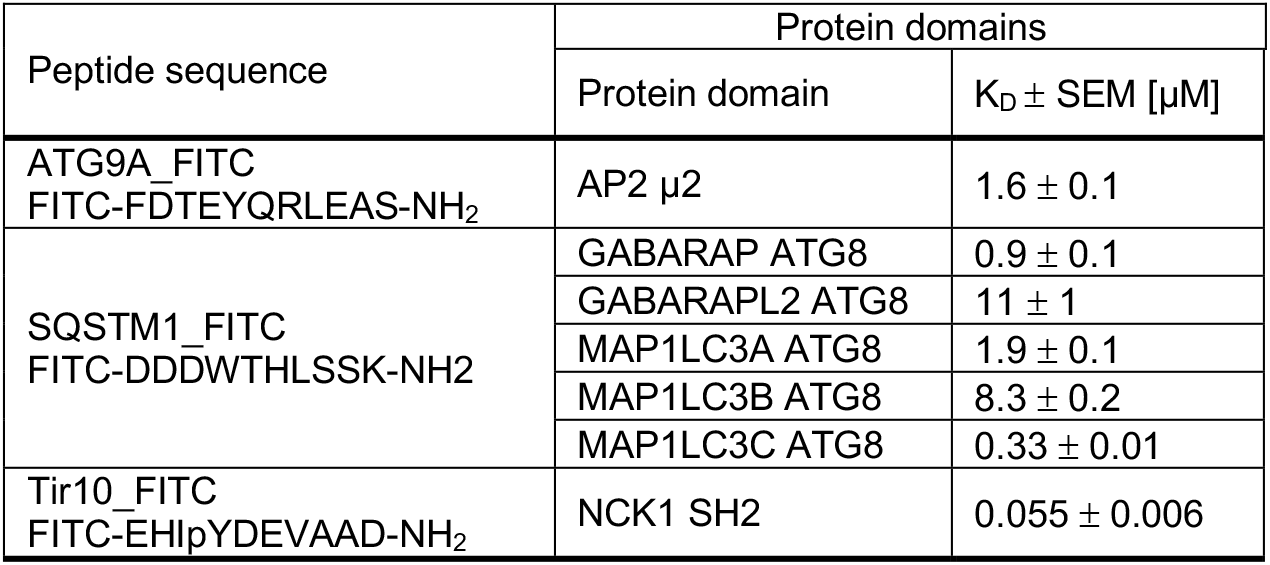
Equilibrium dissociation constant (K_D_) values of AP2 μ2, the ATG8 domains and NCK1 SH2 for the respective FITC-labelled peptides (ATG9A, SQSTM1 and Tir10). Indicated errors are the errors of the mean (SEM).

The NCK1 SH2 did not bind to either of the tested peptides, which may reflect the difficulties in accurately predicting binding motifs of SH2 domains with relatively poorly defined consensus motifs. However, it should be noted that the predicted motif might serve as an interaction site for other SH2 domains. In contrast, we demonstrate a weak interaction between AP2 μ2 and ACE2 peptide with preferential binding to unphosphorylated ACE2 peptide (K_I_ value = 101 ± 5 μM). Phosphorylation at Tyr781 disabled the interaction with AP2 μ2, which is consistent with previous reports (37,38). We thus confirm the presence of a weak endocytic sorting signal in ACE2 C-terminal tail, that may be regulated by phosphorylation.

### The integrin β_3_ tail contains a phospho-regulated LIR motif

After the evaluation of the proposed SLiMs in the ACE2 tail we assessed the predicted LIR motifs in integrin tails, using the integrin β3 tail as a model system. We determined K_I_ values for unphosphorylated peptide as well as phospho-peptides spanning the region of the potential binding motif by displacement experiments. The hydrophobic motif is represented by Phe780 and Ile783 and interspaced by two variable amino acids (Figure 1). Preceding the motif, there is a Ser/Thr-rich sequence (residues 777-779), suggested to be subjected to phosphorylation. In context of the LIR motif, Tyr785 emerges equally as relevant potential phosphosite (14).

Displacement curves were obtained in the same fashion as described above and KI values concomitantly calculated (Figure 4, Table 3). We can in particular demonstrate that the interaction is tuned by phosphorylation of residues upstream and downstream of the core hydrophobic motif. We found that the unphosphorylated peptide is essentially not bound by the ATG8 domains tested. Phosphorylation of Ser778 located upstream of the hydrophobic motif strengthened ATG8 binding for all domains tested, while Thr779 phosphorylation enhanced the affinity for MAP1LC3C, but had no effect on the binding of the other ATG8 domains. Phosphorylation of the downstream residue Tyr785 conferred improved affinity in all tested cases. Concomitant phosphorylation of Thr779 and Tyr785 promoted ATG8 domain binding (double phosphorylated integrin β3 peptide) and increased the binding affinity of MAP1LC3C even further, indicating a synergistic effect of the two phosphosites (Figure 4). We thus clearly demonstrate a phospho-dependent interaction between the ATG8 proteins at the biophysical level.

**Figure 4:**
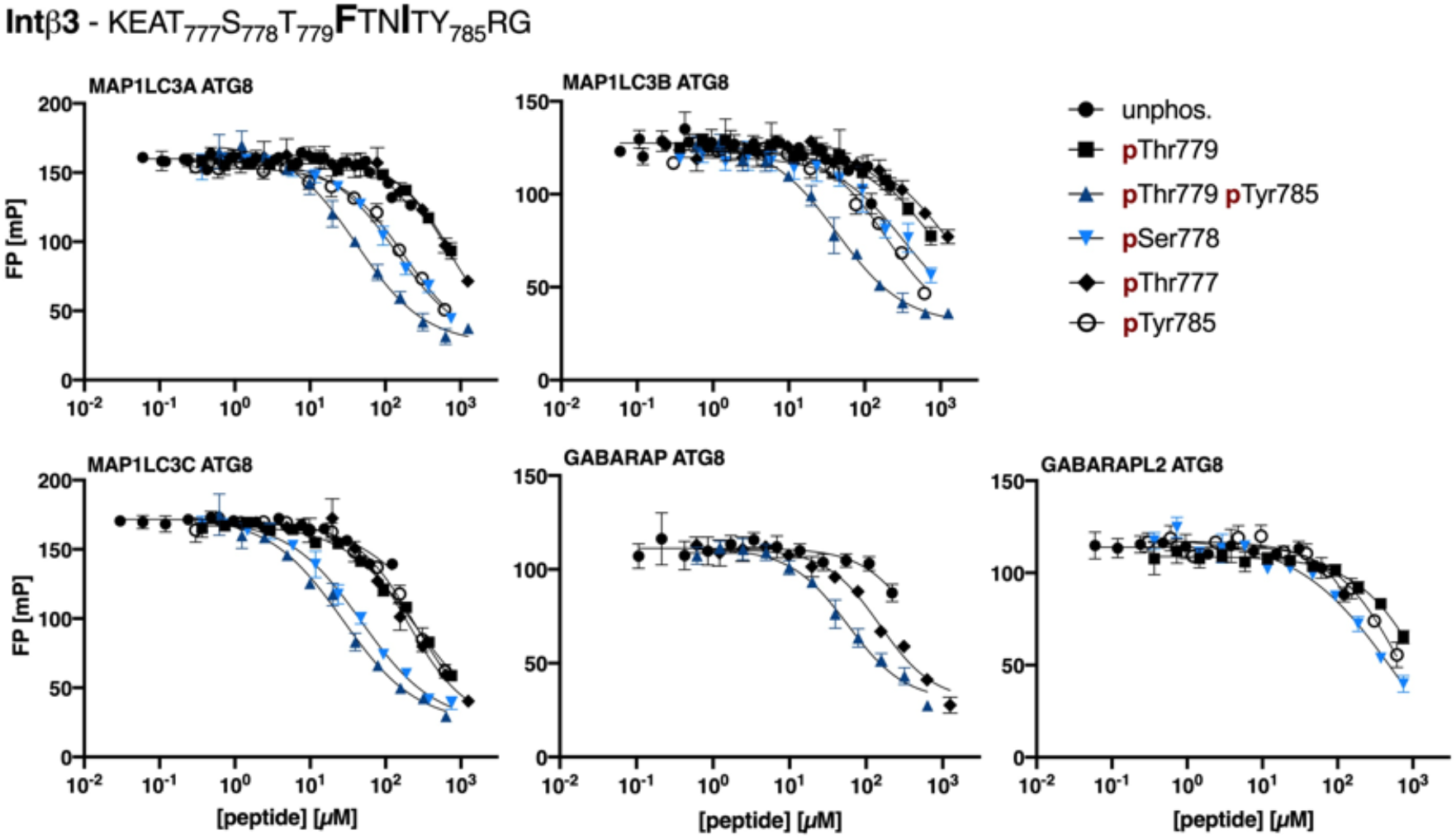
Fluorescence polarisation displacement curves obtained using unphosphorylated and phosphorylated peptides representing the cytoplasmic integrin β3 tail. Peptide sequence and sampled phosphosites are indicated in the figure.

**Table 3:**
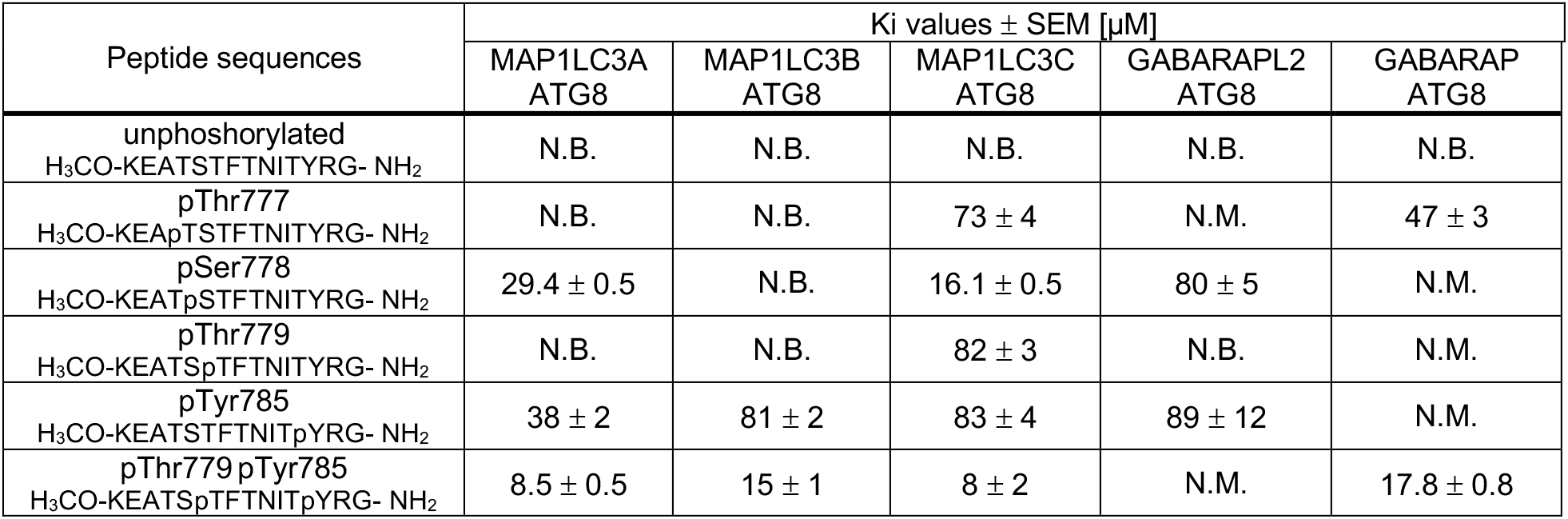
K_I_ values for ATG8 domains calculated from displacement experiments using unphosphorylated integrin β_3_ peptide as well as phosphorylated peptides (pThr777, pSer778, pThr779, pTyr785 or pThr779 pTyr785). Indicated error is the error of the mean (SEM). N.B. indicates no or low affinity (Ki >100 μM). N.M. indicates not measured.

## Discussion

We provide biophysical evidence for the functionality of SLiMs harboured by the cytoplasmic tails of ACE2 and integrin β_3_, providing possible links between the receptors and both endocytosis and autophagy.

### Validation of trafficking motifs in the C-terminal tail of ACE2

Coronaviruses, including SARS-CoV-2, are proposed to utilise either the endocytic or non-endosomal pathway for cell entry (39). Our biophysical validation confirms the presence of an endocytic AP2 μ2 binding motif in the ACE2 C-terminus, which is negatively regulated by Tyr781 phosphorylation. This finding supports the possibility of clathrin-dependent endocytosis of SARS-CoV-2 upon receptor binding. However, it will of course be necessary to assess the biological relevance of the validated ACE2-AP2 μ2, particularly in light of its low affinity (100 μM). Concerning the latter, it can be noted that, while affinities of AP2 core complexes for endocytic sorting signals are reported in nanomolar range, measured K_D_ values of the AP2 μ2 subunit alone vary between 10 to 70 μM (40–42). Those reports contextualise the here reported affinity to a reasonable range.

Through our small-scale sampling, guided by tissue distribution and PDZ binding motif preferences, we demonstrate that SHANK1 PDZ and NHERF3 PDZ1 bind to the C-terminus of ACE2 with similar affinities. Between the two PDZ domains, the SHANK1 interaction appears less physiologically relevant because SHANK proteins have a known role in the structural organisation of the postsynaptic density (43). In contrast, NHERF3 serves as a scaffold protein that regulates the surface expression of plasma membrane proteins in the apical domain of epithelial cells (44,45), which agrees with the fact that ACE2 is found enriched on the apical surface of conducting airway epithelia (46). NHERF3 has moreover been reported to interact with the glutamate transporter excitatory amino acid carrier (47). This transporter harbours, apart from the PDZ binding motif and similar to ACE2, an endocytic sorting signal demonstrated to mediate AP2 μ2-dependent endocytosis.

Hence, it is possible that an equivalent mechanism orchestrates ACE2 surface stabilisation (PDZ-mediated) and internalisation (AP2-mediated). It appears feasible that the interaction between NHERF3 and ACE2 is functionally relevant due to their involvement in similar biological processes and suggested shared tissue distribution. Our analysis, even though limited by the set of PDZ domains tested, points towards a specificity of ACE2-PDZ interactions. However, we note that there are more than 260 human PDZ domains with partially overlapping specificities (32), and that other class I PDZ domains with similar binding preferences, such as the sorting nexin 27 (SNX27) PDZ domain, may also be physiological binders. SNX27, involved in retrograde transport from endosome to plasma membrane and hence in the recycling of internalised transmembrane proteins (48), was recently found as a host factor required for efficient entry of an engineered SARS-CoV-2 variant, the spike protein of which contains a deletion at the S1/S2 subunit cleavage site (49). Thus, it is plausible that SNX27 is involved in recycling ACE2 to the plasma membrane, which would in extension promote viral entry. Alternative methods, such as the hold-up assay against the full PDZ domain collection (50), may shed further light on the PDZ specificity profile of the ACE2 C-terminus in the future. Finally, in terms of the NPxF motif we find that the interaction with SNX17 FERM is the most promising lead, and note that SNX27 is involved in preventing lysosomal degradation of β integrins by binding to a NPxY motif in the cytoplasmic tails of β integrin (35).

### Functional LIR motifs in integrin β3 as potential linkage of SARS-CoV-2 infection to autophagy

We confirm that the ATG8 domains (MAP1LC3A, -B, -C, GABARAP, -L2) bind the integrin β3 cytosolic tail in a phospho-dependent fashion. Phosphorylation upstream of the core LIR motif in the integrin β3 peptide enhanced ATG8 domain binding with a pronounced synergistic effect between the Thr779 and Tyr785 phosphosites. Notably, phospho-regulation of the LIR motif has already been suggested by various reports (51). Future systematic analysis of the effects of phosphosites relative to the core LIR motif will provide additional details of the regulation. The functionality of the LIR binding motif in integrin β_3_ provides a direct link between the receptor and the autophagic machinery, establishing a possible mechanism how SARS-CoV-2 could hijack this pathway for its propagation. This hypothesis appears particularly feasible in light of the fact that ATG8 domains act as adaptors recruiting LIR-containing proteins to phagosomal membranes (51).

How the autophagic machinery is deployed during coronavirus infection is still under debate with the majority of research conducted on viruses such as SARS-CoV and mouse hepatitis virus (39). Recent interest is driven by e.g. the finding that a number of inhibitors with impact on the autophagic flux reduce the cytopathic effect of SARS-CoV-2 infection in Vero-E6 cells (52). Further, a study repurposing approved compounds identified apilimod, a PIKfyve kinase inhibitor (53), as a potent inhibitor of viral replication in an ex vivo lung culture system (54). PIKfyve kinase activity has been linked to critical membrane trafficking events due to the generation of PI(3,5)P2 (55) and its inhibition by apilimod to induce secretory autophagy (56). The data, which we provide here, may hence shed light on the interconnection between infection and the autophagic path. Redistribution of membranes important for viral replication, which might be suggested by apilimod treatment, may sensitively affect viral entry and replication. However, *in-cellulo* follow-up studies are required to test whether the virus deploys the potential protein-protein interactions of integrin β_3_.

From a more general perspective, an inverse relationship between autophagy and integrin-dependent anchorage signalling has been proposed, implying that the autophagic machinery counteracts anoikis (detachment-induced apoptosis) (57). For instance, downregulation of integrin β_3_ resulted in upregulation of autophagy markers (58). In line, it was observed that blockage of α_V_β_3_ signalling results in inhibition of adhesion and concomitant activation of the autophagic machinery, manifested in increased MAP1LC3B expression (59). Conclusive evidence linking integrin signalling and autophagy has thus been gathered, whereas the molecular basis underlying vesicular rearrangements and the trafficking of integrins has not been established so far.

The phospho-dependence of the LIR motif, which we demonstrate here, might explain the conditional regulation of the interaction. Intriguingly, Shc has been identified as a key mediator of anoikis and deletion of its PTB domain to result in loss of anoikis activity (60). Further, binding of the PTB domain to the integrin β_3_ tail is dependent on the phosphorylation of Tyr785 (61,62), whereas double phosphorylation of Thr779 and Tyr785 abolishes the interaction (30). As we could show that the double phosphorylation enables binding of MAP1LC3s and GABARAP synergistically, it can be hypothesised that the integrin β_3_ tail serves as a molecular platform signalling for either the induction of anoikis or protective autophagy.

## Conclusion

To conclude, we report on the functionality of some of the predicted binding motifs harboured by the cytoplasmic tail of ACE2 including a selective class I PDZ binding motif and a weak endocytic sorting signal. The presence of these motifs points to protein-protein interactions downstream of receptor binding, which may mediate internalisation. This might be relevant in context of both normal cellular physiology and SARS-CoV-2 infection. While we cannot confirm the presence of an autophagy LIR motif in the ACE2 tail, we clearly establish a phospho-dependent LIR motif in the integrin β3 tail. This provides the molecular link between integrin β3 and the autophagic pathway. The study opens the way to researching the mechanistic details of the interplay between autophagy and integrin-mediated signalling. It would be interesting to assess whether the remainder of the β-integrin tails contain phosphomodulated LIR motifs as well. At this point, the coupled involvement of both integrins and the autophagic process in SARS-CoV-2 biology still awaits experimental validation but one can hypothesise that our data contributes to the understanding how the virus might deploy autophagosomes for viral propagation. Contextualising the biological relevance of these confirmed SLiM-based interactions appears as the next logical step.

## Material and Methods

### Peptides

Peptides (Table 4) were obtained from GL Biochem (Shanghai, China) or Genecust (Boynes, France). FITC-labelled peptides were dissolved in DMSO, whereas unlabelled peptides were dissolved in 50 mM sodium phosphate buffer pH 7.4. Either 50 mM sodium phosphate buffer pH 7.4 or phosphate-buffered saline (PBS) pH 7.4 with 3 mM dithiothreitol (DTT) were used and supplemented with 0.05 % Tween for diluting labelled peptides during fluorescence polarisation experiments.

**Table 4:**
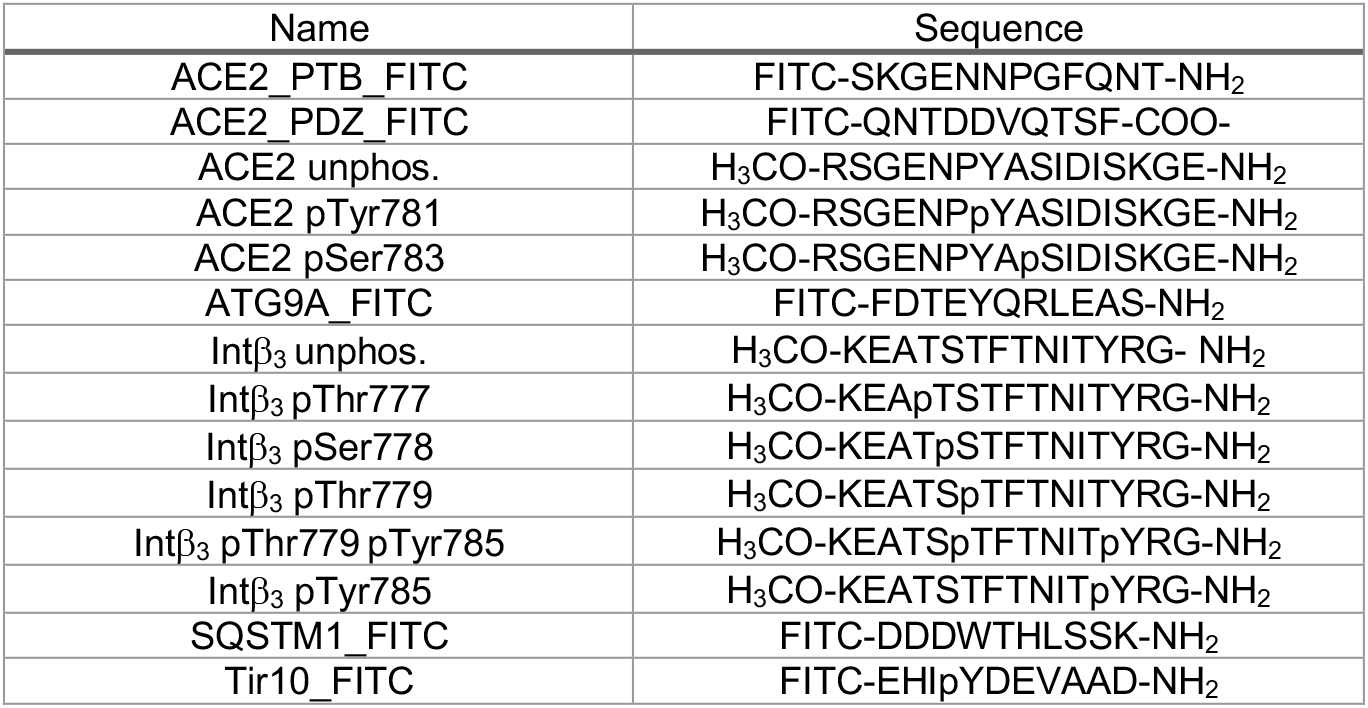
Sequences of peptides used in fluorescence polarisation experiment.

### Protein expression and purification

Protein domains (Table 5) were expressed in E.coli BL21-Gold (DE3), grown in 2YT (10 g/L yeast extract, 16 g/L tryptone and 5 g/L NaCl) and induced at OD600 = 0.7 with 1 mM isopropyl β-D-1-thiogalatactopyranoside followed by 16 h incubation at 18 °C. All domains were glutathione-S-transferase (GST)-tagged, except for the PDZ domains, which were expressed with a maltose binding protein (MBP)-tag. The GST-tagged constructs were purified with glutathione beads (GE Healthcare), whereas ion metal affinity chromatography using Ni^2+^ beads (GE Healthcare) was used for PDZ domains. For affinity measurements, the GST tag was removed by thrombin or tobacco etch virus protease (SNX17 FERM) cleavage, whereas the fluorescence polarisation experiments of the PDZ domains were performed on the tagged protein domains. After purification, protein domains were dialysed in 50 mM sodium phosphate buffer pH 7.4 1 mM DTT or PBS pH 7.4 3 mM DTT (PDZ domains). For the complex in displacement experiments, proteins were supplemented with 0.05 % Tween.

**Table 5:** Protein domains and the encoding plasmids Species is human unless otherwise stated.

| Name | Plasmid |
| --- | --- |
| MAP1LC3A ATG8 (2-119) | pGEX-4T1 |
| MAP1LC3B ATG8 (2-119) | pGEX-4T1 |
| MAP1LC3C ATG8 (2-125) | pGEX-4T1 |
| GABARAPL2 ATG8 (2-115) | pGEX-4T1 |
| NCK1 SH2 (275-377) | pGEX-2T |
| MAGI1 PDZ1 (1-106) | pETM41 |
| NHERF3 PDZ1 (2-107) | pETM41 |
| NHERF3 PDZ2 (132-224) | pETM41 |
| SCRIB PDZ1 (718-814) | pETM41 |
| SHANK1 PDZ (655-761) | pETM41 |
| SNTA1 PDZ (84-171) | pETM41 |
| TAX1B3 PDZ (7-124) | pETM41 |
| DOK1 PTB (151-256) | pH1003 |
| FRS2 PTB (8-136) | pH1003 |
| IRS1 PTB (157-267) | pH1003 |
| TLN1 PTB (442-770) | pH1003 |
| TLN2 PTB (309-401) | pH1003 |
| mouse SNX17 FERM (109-388) | pMCSG10 |

### Fluorescence polarisation experiments

FITC-labelled peptides were used as probes (excitation: 485 nm, emission: 535 nm) and the emitted light was measured in a SpectraMax iD5 Multi-Mode Microplate Reader (Molecular Devices) in order to calculate the mP signal. Experiments were performed at room temperature using non-binding black half area 96-well microplates (Corning^®^) (50 μL/well). Saturation experiments were conducted using 5 nM FITC-labelled peptide and a dilution series of respective protein domain. Displacement curves were obtained by using a premixed complex of protein domain ([protein domain] = 1-2 fold K_D_ of FITC-labelled probe) and FITC-labelled peptide (10 nM) and a dilution series of displacing peptide. All measurements were in technical triplicates, except for MAP1LC3B and GABARAPL2 ATG8 domain binding to ACE2 pTyr785 peptide for which a duplicate of the displacement was recorded. Ki values were calculated as described elsewhere (63).

## Acknowledgement

The MAP1LC3 constructs were kind gifts from Dr. Andreas Ernst (64), the PTB domain constructs were provide by Prof. Sachdev Sidhu (University of Toronto), the PDZ domain constructs were provided by Dr. Renaud Vincentielli & Prof. Carlos Fontes (65), the NCK SH2 domain was a gift from Prof. Bruce Mayer (Addgene plasmid # 46457) and the SNX17 FERM domain construct was provided by Prof. Brett M Collins (33). We thank Dr. Toby Gibson (EMBL Heidelberg) and his team for useful discussions and constructive comments on the manuscript.

## Funding

This work was supported by grants to YI from the Swedish research council (2016-04965) and the Swedish foundation for strategic research (SB16-0039).

## Author contributions

YI and JK conceived experiments. MA and JK performed experiments and analysed results. JK and YI wrote the manuscript. All authors approved the final version.

## Author contributions

The authors declare no competing interests.

## Notes

### Competing Interest Statement

The authors have declared no competing interest.

